# Kir6.2-containing K_ATP_ channels are necessary for glucose dependent increases in amyloid-beta and Alzheimer’s-related pathology

**DOI:** 10.1101/2022.02.20.481215

**Authors:** John Grizzanti, William R. Moritz, Morgan C. Pait, Molly Stanley, Sarah D. Kaye, Caitlin M. Carroll, Nicholas J. Constantino, Lily J. Deitelzweig, Noelle Nicol, James A. Snipes, Derek Kellar, Emily E. Caesar, Jasmeen Dhillon, Maria S. Remedi, Celeste M. Karch, Colin G. Nichols, David M. Holtzman, Shannon L. Macauley

## Abstract

Increased neuronal excitability contributes to amyloid-β (Aβ) production and aggregation in the Alzheimer’s disease (AD) brain. Previous work from our lab demonstrated that hyperglycemia, or elevated blood glucose levels, increased brain excitability and Aβ release potentially through inward rectifying, ATP-sensitive potassium (K_ATP_) channels. K_ATP_ channels are present on several different cell types and help to maintain excitatory thresholds throughout the brain. K_ATP_ channels are sensitive to changes in the metabolic environment, which are coupled to changes in cellular excitability. Therefore, we hypothesized that neuronal K_ATP_ channels are necessary for the hyperglycemic-dependent increases in extracellular Aβ and eliminating K_ATP_ channel activity will uncouple the relationship between metabolism, excitability, and Aβ pathology. First, we demonstrate that Kir6.2/*KCNJ11*, the pore forming subunits, and SUR1/*ABCC8*, the sulfonylurea receptors, are predominantly expressed on excitatory and inhibitory neurons in the human brain and that cortical expression of *KCNJ11* and *ABCC8* change with AD pathology in humans and rodent models. Next, we crossed APP/PS1 mice with Kir6.2 -/- mice, which lack neuronal K_ATP_ channel activity, to define the relationship between K_ATP_ channels, Aβ, and hyperglycemia. Using *in vivo* microdialysis and hyperglycemic clamps, we explored how acute elevations in peripheral blood glucose levels impacted hippocampal interstitial fluid (ISF) glucose, lactate, and Aβ levels in APP/PS1 mice with or without K_ATP_ channels. Kir6.2+/+, APP/PS1 mice and Kir6.2-/-, APP/PS1 mice were exposed to a high sucrose diet for 6 months to determine the effects of chronic hyperglycemia on Aβ deposition. We found that elevations in blood glucose levels correlate with increased ISF Aβ, amyloidogenic processing of amyloid precursor protein (APP), and amyloid plaque pathology in APP/PS mice with intact K_ATP_ channels. However, neither acute hyperglycemia nor chronic sucrose overconsumption raised ISF Aβ or increased Aβ plaque burden in APP/PS1 mice lacking Kir6.2-K_ATP_ channel activity. Mechanistic studies demonstrate ISF glucose not only correlates with ISF Aβ but also ISF lactate. Without K_ATP_ channel activity, ISF lactate does not increase during hyperglycemia, which correlates with decreased monocarboxylate transporter 4 (MCT4) expression, a lactate transporter responsible for astrocytic lactate release. This suggests that K_ATP_ channel activity regulates ISF lactate during hyperglycemia, which is important for Aβ release and aggregation. These studies identify a new role for Kir6.2-K_ATP_ channels in Alzheimer’s disease pathology and suggest that pharmacological antagonism of Kir6.2-K_ATP_ channels holds therapeutic promise in reducing Aβ pathology, especially in diabetic and prediabetic patients.

## INTRODUCTION

Epidemiological studies demonstrate that patients with type-2-diabetes (T2D) have a 2-4 fold increased risk for developing Alzheimer’s disease (AD)[1-4]. Although insulin dysregulation certainly plays a role in the progression of T2D and AD, simple questions regarding how alterations in peripheral glucose metabolism affect brain function, cellular excitability, and proteostasis in AD remain relatively unexplored. Since T2D patients suffer from chronic hyperglycemia, or elevated blood glucose levels, abnormal glucose homeostasis likely plays a role in the development and propagation of AD-related pathology. In fact, recent studies demonstrate that chronic hyperglycemia increases dementia risk [5] and individuals with elevated blood glucose levels have a higher rate of conversion from mild cognitive impairment to AD [6]. Chronic hyperglycemia is also linked to alterations in brain spontaneous activity, decreased functional connectivity, and increased neuronal loss [7], suggesting that alterations in cerebral metabolism are linked to aberrant neuronal activity. This is particularly noteworthy since brain hyperexcitability may drive Aβ/tau accumulation as increased neuronal activity has been shown to stimulate Aβ/tau release, propagation, and aggregation [8-19]. Reciprocally, the formation of amyloid plaques and neurofibrillary tangles feeds forward to cause brain hyperexcitability[19-23]. Therefore, risk factors that alter neuronal excitability can have a significant impact on Aβ/tau metabolism and AD pathogenesis.

To understand how changes in peripheral metabolism impact Aβ pathology, our initial work investigated whether hyperglycemia, or elevated blood glucose levels, or hyperinsulinemia, elevated blood insulin levels, increased amyloid-β (Aβ) levels in the brain’s interstitial fluid (ISF)[12, 24]. By coupling *in vivo* microdialysis and glucose clamps, we developed a novel approach to dynamically modulate systemic blood glucose and/or insulin levels while sampling proteins and metabolites within the brain’s ISF in unanesthetized, freely moving mice. We found that hyperglycemia increased Aβ production in the hippocampus through an activity dependent mechanism; an effect that is exacerbated in mice with amyloid plaques. Interestingly, hyperinsulinemia did not have the same effect, suggesting hyperglycemia is a more potent driver of Aβ production than hyperinsulinemia. We also found a direct correlation between ISF glucose and ISF Aβ concentrations in our acute rodent studies as well as in vervet monkeys with type-2-diabetes where blood glucose levels correlated with decreased CSF Aβ42, a biomarker of Alzheimer’s disease pathology [25]. Interestingly, our rodent studies also found that hyperglycemia increased ISF lactate, a marker of neuronal activity [12, 26], suggesting alterations in cerebral metabolism are tied to brain excitability and Aβ release.

Using a pharmacological approach, we identified inward rectifying, ATP-sensitive potassium (KATP) channels as a possible mechanistic link between elevated glucose levels, neuronal excitability, and Aβ metabolism[12]. K_ATP_ channels are found on the plasma membranes of excitable cells and link changes in metabolism with cellular excitability. Composed of four pore-forming (Kir6.1, Kir6.2) subunits and four sulfonylurea receptor (SUR1, SUR2A, SUR2B) binding sites, K_ATP_ channels play a role in a variety of physiological and pathological conditions[27, 28]. In the pancreatic beta cell, rising blood glucose levels increase intracellular ATP levels, trigger K_ATP_ channel closure, depolarize cellular membranes, and cause insulin secretion. In the cardiovascular system, K_ATP_ channels regulate vasodilation, vasoconstriction, and vascular tone. In the brain, K_ATP_ channels are found on both neurons and glia, where increased glucose metabolism causes neuronal K_ATP_ channel closure, membrane depolarization, and increased cellular excitability [29, 30]. Thus, normal physiological function in a wide range of tissues is closely tied to energy availability and K_ATP_ channel activity.

Therefore, the goal of this study was to explore the role of KATP channels in AD. First, we used publicly available databases from human post-mortem RNA-seq studies to assess determine the cell-type specific expression of Kir6.2 (e.g. *KCNJ11)* and SUR1 (e.g. *ABCC8*) and how it changed across the AD continuum. Next, we used a genetic approach to ablate KATP channel activity in APP/PS1 mice to determine 1) whether KATP channels are necessary for hyperglycemia-dependent increases in ISF Aβ and neuronal activity and 2) whether chronic sucrose overconsumption increases Aβ-related pathology in a KATP channel dependent manner. Deletion of the KATP channel subunit, Kir6.2 (Kir6.2-/-), results in ablation of channel activity in neurons and causes cells to be glucose insensitive. Kir6.2-/- mice are viable, suffer transient hyperinsulinemia as neonates and mild glucose intolerance as adults, but otherwise thrive into adulthood [31, 32]. Thus, we crossed APPswe/PSEN1dE9 (e.g. APP/PS1 mice) to mice lacking neuronal KATP channels, the Kir6.2 -/- mice. Using in vivo microdialysis and hyperglycemic clamps, we demonstrated that doubling peripheral blood glucose levels increases hippocampal ISF glucose, lactate, and Aβ levels in APP/PS1 mice with functioning KATP channels (e.g. Kir6.2+/+, APP/PS1 mice). Chronically, we exposed Kir6.2+/+, APP/PS1 mice and Kir6.2-/-, APP/PS1 mice to high sucrose water for 6 months and demonstrated that chronic sucrose overconsumption increased both the amyloidogenic processing of APP and Aβ deposition in APP/PS1 mice with KATP channels. However, in APP/PS1 mice lacking Kir6.2-KATP channel activity (e.g. Kir6.2-/-, APP/PS1 mice), neither acute hyperglycemia nor chronic high sucrose water increases ISF Aβ or Aβ plaque burden, suggesting Kir6.2-containing KATP channels are necessary for hyperglycemia dependent increases in Aβ-related pathology. The relationship between glucose and Aβ appears to be mediated by changes in lactate production and release, which is also modulated by KATP channel activity. These data suggest that neuronal KATP channels are necessary and sufficient for coupling changes in cerebral metabolism with neuronal activity and Aβ pathology. Furthermore, this work suggests that pharmacological antagonism of Kir6.2-KATP channels holds therapeutic benefit in reducing Aβ pathology for diabetic and prediabetic patients.

## METHODS

### Gene expression alternations in AD brains

To determine whether K_ATP_ channel genes were differentially expressed in AD brain tissue, we analyzed gene expression of *KCNJ11* and *ABCC8* in two publicly available datasets. First, we analyzed gene expression data from the Mayo Clinic Brain Bank (Mayo) RNAseq study from the Accelerating Medicines Partnership – Alzheimer’s Disease (AMP-AD) portal: the temporal cortex of 80 control, 82 AD, and 29 Pathologic Aging (PA) brains (syn3163039; syn5550404). Multi-variable linear regression analyses were performed using CQN normalized gene expression measures and including age at death, gender, RNA integrity number, brain tissue source, and flowcell as biological and technical covariates [33]. Second, we used Mathys et al. (synapse.org: syn18485175) [34] single nuclei transcriptomics (snRNA-seq) analysis of AD brains to determine the cell type specific expression of *KCNJ11* and *ABCC8* in post-mortem human prefrontal cortex across the AD continuum (n=48) from the Religious Order Study (ROS) or the Rush Memory and Aging Project (MAP) study[35]. Individuals had either “no-pathology” (N=24) or high levels of Aβ-pathology and other AD-associated pathological changes (N=24). In the Aβ-positive group, the participants were further characterized as “early pathology” suggesting a milder presentation of disease or “late pathology” suggesting a more advanced stage of disease. Individuals were balanced between sexes, age, and years of education. 80,660 droplet-based single-nucleus RNA-seq (snRNA-seq) profiles were analyzed and differential expression assessed. Extensive description of methodology used is described here [34]. Data was filtered per Mathys et al. and clustered using the SCANPY package (Wolf et al. 2018) in Python. The Leiden algorithm was applied to identify cell clusters and a Uniform Manifold Approximation and Projection (UMAP) algorithm was used for dimension reduction and visualization of clustering results[36].

### APPswe/PS1ΔE9 and Kir6.2-/- mice

APPswe/PS1ΔE9 mice on a B6C3 background (APP/PS1; Jankowsky et al, Hum Mol Genet, 2004) were crossed to Kir6.2-/- (Miki et al, PNAS, 1998) or Kir6.1-/- [37] for these studies. To generate APP/PS1 mice homozygous knockout (-/-) for Kir6.2, we bred Kir6.2_-/-_ mice (gift from Dr. Colin Nichols) with APP/PS1 mice, generating Kir6.2+/+, APP/PS1 and Kir6.2-/-, APP/PS1 mice for the acute and chronic experiments. For reverse microdialysis experiments, we also generated mice homozygous knockout (-/-) for Kir6.1 (gift from Dr. Colin Nichols) crossed to APP/PS1 mice, generating Kir6.1-/-, APP/PS1 mice. Mice were given food and water ad libitum and maintained on a 12:12 light/dark cycle. All procedures were carried out in accordance with an approved IACUC protocol from Washington University School of Medicine or Wake Forest School of Medicine.

### In vivo microdialysis and hyperglycemic clamps

Five-days prior to glucose clamps [38], 3-month Kir6.2+/+, APP/PS1 and Kir6.2-/-, APP/PS1 mice (n=7- 8 mice/group) were anesthetized via isoflurane and tapered catheters (MRE025 tubing, Braintree Scientific) inserted into the jugular vein and the femoral artery, and sutured into place. The catheter lines were filled with polyvynilporolidone (PVP), the ends double knotted, and a suture affixed to the ends. A small incision was made between the scapulae and the lines tunneled to this area for externalization prior to glucose clamps. Two days prior to glucose clamps, guide cannulas (BR-style, BIoanalytical Systems) were stereotaxically implanted into the hippocampus (from bregma, A/P: -3.1mm, M/L: -2.5mm, D/V: -1.2mm at 12° angle) and secured into place with dental cement. Cresyl violet staining was used to validate probe placement post-mortem. One day prior to glucose clamps, the mice were transferred to Raturn sampling cages (Bioanalytical Systems) and microdialysis probes (2 mm; 38 kDa molecular weight cut-off; BR-style, Bioanalytical Systems) inserted into the guide cannula, connected to a syringe pump and infused with artificial cerebrospinal fluid (aCSF; 1.3mM CaCl_2_, 1.2mM MgSO_4_, 3mM KCl, 0.4mM KH_2_PO_4_, 25mM NaHCO_3_ and 122mM NaCl; pH=7.35) at a flow rate of 1 μl/minute. At this time, the jugular and femoral lines were externalized and the PVP removed. The femoral and jugular lines were flushed, connected to a syringe pump and slowly infused with 0.9% sodium chloride at a flow rate of 1μl/min overnight to prevent clots. Hourly collection of hippocampal ISF began. The following morning, mice were fasted 4-5 hours prior to and during the 4-hour glucose clamps. For the duration of the clamp, the jugular vein was infused with a 12.5% dextrose solution in PBS (control mice received PBS alone) at a variable flow rate. Every 10 minutes, blood was sampled via the femoral artery and blood glucose concentration assessed using a handheld glucometer (Contour, Bayer). The concentration of blood glucose was targeted to 150-200 mg/dL and the flow rate of the dextrose solution was adjusted accordingly. After the 4-hour clamp, the dextrose solution was stopped, the lines flushed, euglycemia restored, and food returned to the bowls. Hourly ISF collection continued for the duration of the clamp and for 10-15 hours post-clamp.

### Glucose tolerance test

Glucose tolerance test was performed as previously described [15]. Briefly, mice were fasted for 4 h and 2.0 g/kg glucose was administered via i.p injection. Blood samples were taken from tail veins and blood glucose was measured at baseline, 15-, 30-, 45-, 60-, 90-, and 120 minutes from glucose injection using a handheld glucometer (Bound Tree Medical Precision XTRA Glucometer; Fisher).

### Glibenclamide administration via reverse microdialysis

Guide cannula implantation and in vivo microdialysis were performed as described above. Glibenclamide (100μM; Sigma-Aldrich) was infused directly into the hippocampus of Kir6.2+/+, APP/PS1 mice, Kir6.2-/-, APP/PS1 mice, and Kir6.1-/-, APP/PS1 mice (n=4-7) via reverse microdialysis for 3 hours. Changes in ISF glucose, lactate, and Aβ were quantified as described below. Statistical significance was determined using a one-way ANOVA and Dunnett’s multiple comparison post-hoc test. Data is represented by means +/- SEM.

### Aβ_1-X_ ELISA

ISF samples from Kir6.2+/+, APP/PS1 mice, Kir6.2-/-, APP/PS1 mice, and Kir6.1-/-, APP/PS1 mice (n=7- 8/group) collected from in vivo microdialysis experiments were analyzed for Aβ_1-x_ using sandwich ELISAs as previously described[14, 39]. Briefly, Aβ_1-x_ was quantified using a monoclonal capture antibody targeted against Aβ13-28 (m266) and a biotinylated detection antibody targeted against Aβ1-5 (3D6), both generous gifts from Dr. Ron DeMattos, Eli Lilly and Co., Indianapolis, IN. After incubation with streptavidin-poly-HRP-20, the assay was developed using Super Slow TMB (Sigma) and the plates read on a Bio-Tek Synergy 2 plate reader at 650nm. Statistical significance was determined using a two-tailed, unpaired t-test. Data is represented by means +/- SEM.

### Chronic high sucrose diet

Kir6.2+/+, APP/PS1 mice and Kir6.2-/-, APP/PS1 mice were randomized to either regular drinking water group or a high sucrose (104mM glucose, 128mM fructose) drinking water group (n=8-11 mice/group) at 3 months of age. Mice were subject to either condition for 6 months, where body weight and blood glucose were monitored at baseline, 6, and 9 months of age in all animals. At 9 months of age, mice were sacrificed and processed for several biochemical, metabolic, and AD-related pathology analyses.

### Tissue Collection

At the completion of each experiment, mice were deeply anesthetized using isoflurane, a cardiac puncture to collect blood, and transcardially perfused with heparinized 1X DPBS. Each brain was bisected into left and right hemispheres. One hemisphere was dissected to isolate specific brain regions for biochemical and molecular biology assays and the other hemisphere was fixed in 4% paraformaldehyde (PFA) in 1X DPBS for 48 hours at 4°C. PFA fixed hemispheres were then transferred to 30% sucrose in 1X DPBS for cryoprotection and stored at 4°C until further tissue processing. Brains were sectioned on a freezing microtome at 50um sections.

Blood collected from cardiac puncture during sacrifice was transferred to a microcentrifuge tube coated in EDTA to prevent coagulation. Blood was spun down at 2,000 x g for 10 minutes at 4°C to isolate plasma from the whole blood. The plasma supernatant was then removed, transferred to a clean tube, and stored at - 80°C until use.

### Insulin ELISA

Plasma insulin levels of 9-month-old animals were measured via mouse ultra-sensitive insulin ELISA (ALPCO,) according to manufacturer’s protocol. Briefly, manufacturer provided standard curve samples (0, 0.188, 0.5, 1.25, 3.75, and 6.9 ng/mL) were loaded in triplicate. Duplicate 5 μL samples of plasma were loaded into a 96-well plate that was precoated with a capture insulin antibody. 75 μL conjugated substrate was added to each well and mixed for 2 hours on an orbital shaker at 450 rpms. Wells were then washed 6 times with 1X wash buffer. 100 μL of TMB substrate was then added to each well and mixed for 30 minutes at room temperature. 100 μL of stop solution was added to each well and gently mixed prior to reading. The plate was then immediately read at 450 nm. The standard curve was constructed using a 5-parameter logistic curve and sample optical densities were compared to the standard curve to determine the plasma insulin concentration. Plasma insulin levels were analyzed via one-way ANOVA and Tukey post-hoc tests.

### Glucose and Lactate Measures

Glucose and lactate measurements from blood and hippocampal ISF were quantified using an YSI 2900 analyzer (YSI incorporated) using glucose- and lactate-oxidase method per the manufacturer’s instructions as previously described (Macauley et al 2015 JC). The ratio of plasma glucose to lactate was also calculated by [plasma glucose/plasma lactate]. Plasma levels of glucose, lactate, and glucose: lactate was analyzed via one-way ANOVA and Tukey posthoc tests. Significance was determined at p<0.05, while a trend was p<0.01. Data is represented by means +/- SEM.

### Immunohistochemical staining of brains using anti-Aβ antibodies

Serial sections (300μm apart) through the anterior-posterior aspect of the hippocampus were immunostained for Aβ deposition (anti-HJ3.4B, a generous from David Holtzman). Free floating sections were stained using a biotinylated, HJ3.4 antibody (anti-Aβ1–13, mouse monoclonal antibody) and developed using a Vectastain ABC Elite kit and DAB reaction. Stained brain sections were imaged using a NanoZoomer slide scanner (Hamamatsu Photonics) and the percent area occupied by HJ3.4 was quantified by a blinded researcher as previously described (5, 6). Statistical significance was determined using a two-tailed, unpaired t-test.

### Western Blotting

Western blot analysis was used to measure levels of APP and the c-terminal fragments (CTFs), CTF-α and CTF-β produced from cleavage of APP by α-secretases or β and γ-secretases, respectively (SOURCE). Briefly, hippocampi were homogenized in 1X cell lysis buffer (Cell Signaling) supplemented with protease inhibitor cocktail (Roche), 1mM PMSF (Cell Signaling, MA), 1mM DTT (Sigma-Aldrich), and a phosphatase inhibitor cocktail (Millipore) using a probe sonicator at 30% amplitude, 1 sec pulse with a 5 sec delay 5 times while on ice. Tissue homogenates were then spun down at 10,000 x g for 10 minutes at 4°C and the supernatant was removed and used for immunoblotting. Protein concentrations were analyzed using BCA protein assay kit (Pierce). 15 μg of protein was ran via SDS-PAGE in 10% Tris-tricine gels using BioRad protean mini and then rapid transferred to PVDF membranes using BioRad Semi-dry (BioRad). Membranes were subsequently blocked using 5% BSA in 1X TBST for one hour and then incubated with either 1° (1:1,000) APP and CTFs (Life Technologies) or the loading control (1:50,000) β-actin (Millipore) overnight at 4°C. 2° antibody conjugated with HRP specific to either (1:3,000) goat anti-rabbit (Cell Signaling) or (1:3,000) goat anti-rabbit (Cell Signaling) for 1 hour at room temperature. Chemiluminescence was measured using ECL (EMD Millipore). All images were quantified using ImageJ, normalized to β-actin, and analyzed via one-way ANOVA using Graphpad Prism.

### RNA extraction

Frozen cortex was homogenized in Qiazol (Qiagen) using a sterile, nuclease free hand-held homogenizer (Bel-art) for ∼30 seconds on ice. Samples then sat at room temperature for 5 minutes and then mixed vigorously with chloroform at 1:5 ratio (chloroform: Qiazol) and allowed to sit for 10 minutes at room temperature. Sample mixtures were then spun down at 12,000 x g for 10 minutes at 4 DC. The aqueous layer of RNA was then carefully removed, gently mixed with 1:1 70% EtOH: original volume of Qiazol, and run through a RNeasy spin column (Qiagen) according to manufacturer’s protocol.

### Quantitative Real-Time Polymerase Chain Reaction (qPCR)

1 μg of RNA was then converted to cDNA using High-Capacity cDNA Reverse Transcription kit (Applied Biosystems) according to manufacturer’s protocol. cDNA was then used to run qPCR for several target genes: Kcnj8 (Mm00434620_m1, Life Technologies), Kcnj11 (Mm04243639_s1, Life Technologies), Abcc8(Mm00803450_m1, Life Technologies), Abcc9 (313340540, IDT), Ldha (Mm01612132_g1, Life Technologies), Ldhb (Mm01267402_m1, Life Technologies), Glut1 (316054177, IDT), Glut3 (Mm00441483_m1), Mct2 (Mm00441442_m1, Life Technologies), and Mct4 (Mm0046102_m1, Life Technologies) using Rn-18s (Mm03928990_g1, Life Technologies) as a loading control. Briefly, 20 ng of cDNA was loaded per well with Taqman Fast Advanced Master Mix (Applied Biosystems) along with 0.5 μL of primer for a specific gene of interest and 0.5 μL of Rn-18s primer. qPCR was run using Quant Studio 6 Pro (Applied Biosystems) according to manufactures’ protocols for Taqman primers (Life Technologies) and Taqman Fast Advanced Master Mix. Relative quantities of gene expression were quantified as described in (Livak and Schmittgen, 2001) and gene expression was statistically analyzed via one-way ANOVA using GraphPad Prism.

## RESULTS

### K_ATP_ channels, composed Kir6.2 and Sur1, are found on neurons and have altered expression in the human AD brain and rodent models of AD-related pathology

*KCNJ11*, encodes the pore forming subunit, Kir6.2, and *ABCC8*, encodes sulfonylurea binding site, Sur1. Kir6.2 and Sur1 heterodimerize to form ATP sensitive, inward rectifying potassium (KATP) channels (Figure 1A). To determine whether K_ATP_ channel genes were differentially expressed in AD brain tissue, we analyzed gene expression of *KCNJ11* and *ABCC8* in two publicly available datasets. First, we analyzed gene expression data from the temporal cortex of normal control (NC; n=80), Pathologic Aging (PA=amyloid+; n=29), and Alzheimer’s disease (AD=amyloid+,tau+; n=82) brains in the Mayo Clinic Brain Bank (Mayo) RNAseq study from the Accelerating Medicines Partnership – Alzheimer’s Disease (AMP-AD) portal (Figure 1B). In Pathologic Aging (amyloid+), *KCNJ11* expression was upregulated (beta=0.3641, p val<4.306E-03), while both *KCNJ11* and *ABCC8* expression decreased in the temporal cortex of Alzheimer’s disease (amyloid+tau+) brains (beta=-0.2150, p val<7.294E-02 and beta=-0.2669, p val<3.704E-02, respectively). Next, we leveraged a publicly available single nuclei RNAseq (snRNA-seq) database from post-mortem human prefrontal cortex [34] to explore the cellular localization and expression profile of *KCNJ11* and *ABCC8* across the AD continuum (Figure 1D). First, we found that *KCNJ11* is predominately localized to excitatory neurons (82.47%) or inhibitory neurons (13.66%), and to a lesser degree, oligodendrocyte progenitor cells (1.93%), oligodendrocytes (1.03%), astrocytes (0.58%), or microglia (0.32%; Figure 1E). Similarly, we found that *ABCC8* is predominately localized to excitatory neurons (82.18%) or inhibitory neurons (11.09%), and to a lesser degree oligodendrocyte progenitor cells (4.91%), astrocytes (1.11%), oligodendrocytes (0.62%), microglia (0.05%), endothelial cells (0.03%), and pericytes (0.01%; Figure 1E).When investigating *KCNJ11* expression across the AD continuum, *KCNJ11* expression increases during early AD and late AD in excitatory neurons, while *KCNJ11* expression in inhibitory neurons increases during a late stage of AD (Figure 1E). There is no change in *KCNJ11* expression in glia across the AD continuum (data not shown). Conversely, *ABCC8* expression decreases in both excitatory and inhibitory neurons in early AD and late AD compared to individuals with no pathology (Figure 1E). Lastly, we performed qPCR on the cortex of 9-month-old APP/PS1 and wildtype controls (Figure 1G). *Kcnj11* and *Abcc8* expression is decreased in APP/PS1 mice compared to wildtype controls (Figure 1H). Together, this demonstrate that Kir6.2-K_ATP_ channels are localized to both excitatory and inhibitory neurons and that *KCNJ11* and *ABCC8* expression is altered across the AD continuum in humans and mice.

**Figure 1.**
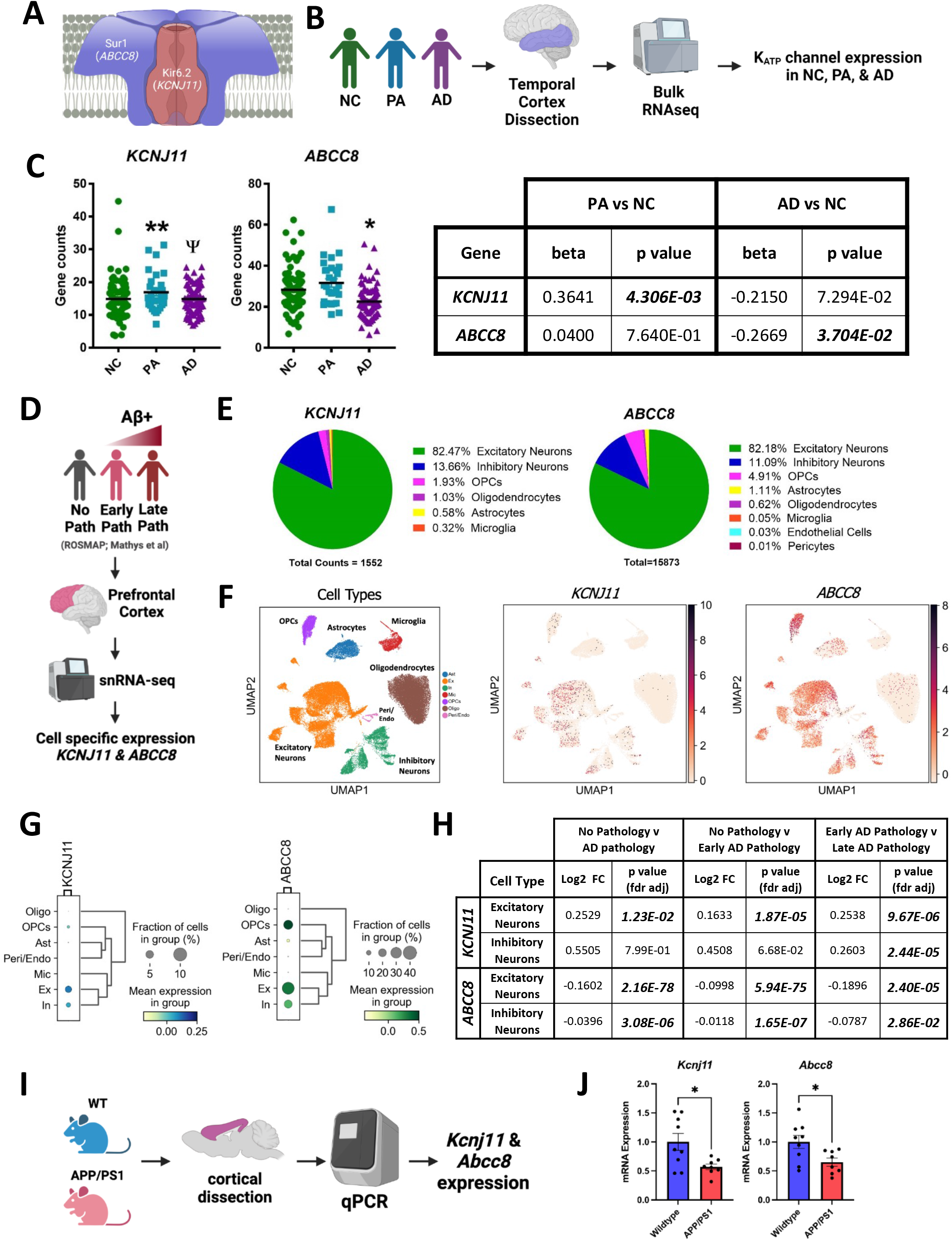
KATP channel expression in post mortem human and mouse brains across the Alzheimer’s disease continuum. (A) ATP-sensitive, inward rectifying potassium (KATP) channels are heteroctameric, composed of 4 pore forming subunits (Kir6.2*/KCNJ11)* and 4 sulfonylurea binding site s(Sur1*/ABCC8)* subunits. (B) Workflow to explore how K_ATP_ channel genes (*KCNJ11, ABCC8*) in the temporal cortex change due to AD-related pathology using the Mayo RNAseq database. (C) *KCNJ11* expression increases in pathological aging (PA; amyloid+) and trends towards a decrease in Alzheimer’s disease (AD; amyloid+,tau+). Similarly, ABCC8 trends higher in PA and is significantly reduced in AD. (D) Workflow to explore cell type specific changes in *KCNJ11* and *ABCC8* expression in post-mortem human brains using single nuclei RNAseq (snRNA-seq) database generated by Mathys et al (2019). (E) *KCNJ11* and *ABCC8* expression is largely found on excitatory and inhibitory neurons (>96%), but also localized to glia. (F) NC and AD samples were integrated into a single dataset and clustered into cell types. UMAP representation of different CNS cell types, including relative expression for *KCNJ11* and *ABCC8*. (G) Gene expression dot blot for *KCNJ11* and *ABCC8* demonstrating relative expression levels in each cell type. (H) Comparing post mortem brains with no AD pathology (n=24), early AD pathology (n=15), and late AD pathology(n=9), *KCNJ11* expression is increased on excitatory neurons with early and late pathology while *KCNJ11* expression is increased on inhibitory neurons at a late stage of disease. (I) Workflow for exploring *Kcnj11* and *Abcc8* expression in the mouse cortex of APP/PS1 and WT mice. (J) Decreased *Kcnj11* and *Abcc8* expression in 9 month APP/PS1 cortex compared to 9 month WT mice.

### Kir6.2-KATP channels link acute changes in brain glucose with ISF Aβ and lactate levels

To explore the role of Kir6.2- K_ATP_ channels in Alzheimer’s-related pathology, we crossed mice deficient in Kir6.2 (e.g. Kir6.2-/- mice) [31] with APPswe/PSEN1dE9 (e.g. APP/PS1) [40] mice to generate mice overexpressing Aβ that were also deficient in neuronal K_ATP_ channel activity (e.g. Kir6.2-/-, APP/PS1). This model can then be used to explore whether neuronal K_ATP_ channels link changes in cerebral metabolism with neuronal excitability and Aβ release. We previously combined hyperglycemic clamps and in vivo microdialysis as a tool to investigate how dynamic changes in blood glucose levels alter metabolites and proteins in real-time in the brain’s ISF of unrestrained, freely moving mice [12, 41]. By coupling this approach with our newly developed mouse model (e.g. Kir6.2-/-, APP/PS1), we were able to explore whether neuronal K_ATP_ channels are necessary for glucose-dependent increases in hippocampal ISF Aβ and ISF lactate (Figure 2A).

**Figure 2.**
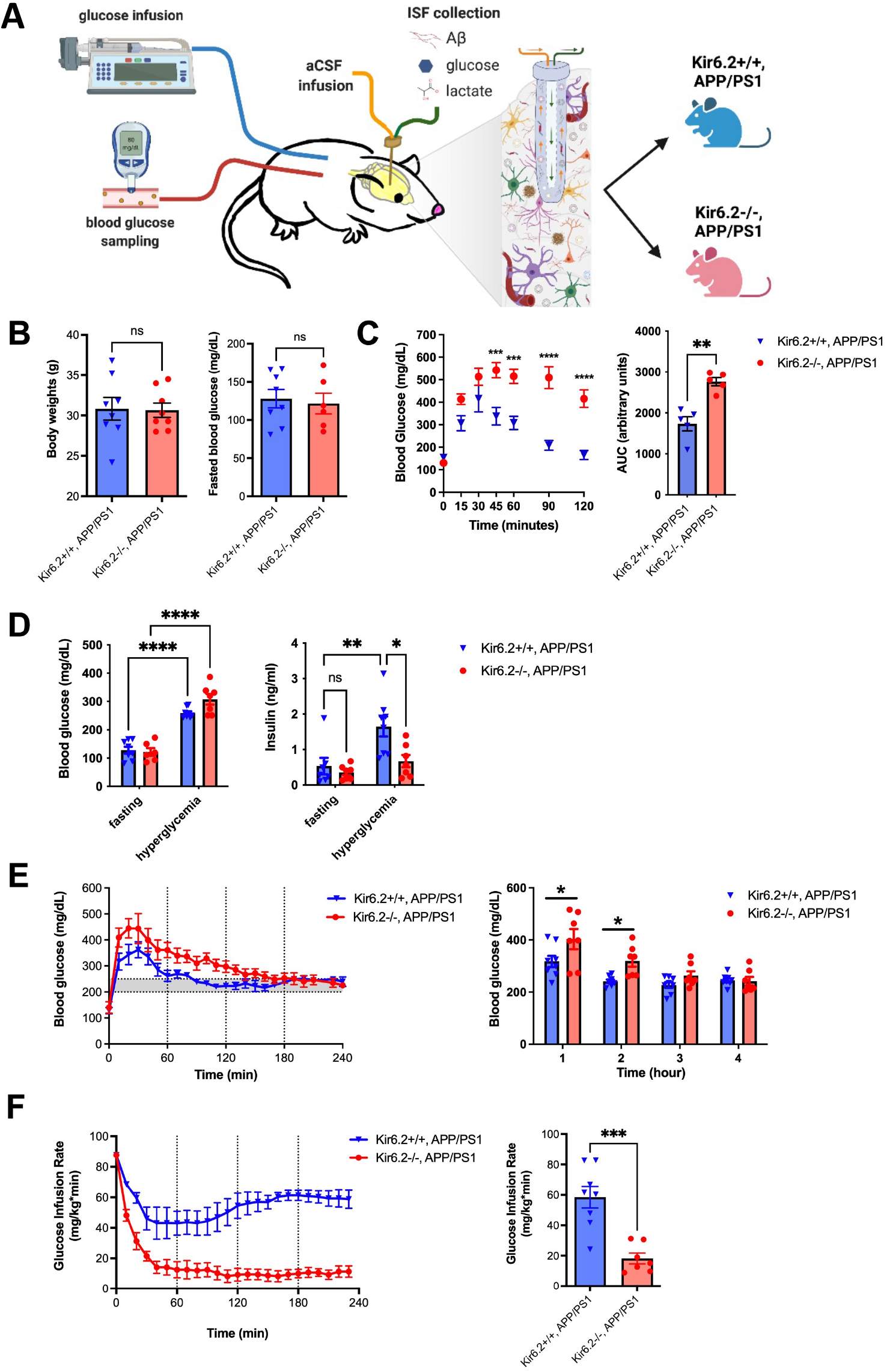
The effects of hyperglycemic clamps on blood glucose, serum insulin, and glucose infusion rate in Kir6.2-/-, APP/PS1 mice compared to in Kir6.2+/+, APP/PS1 mice. (A) Schematic of experimental approach where hyperglycemic clamps are paired with in vivo microdialysis to assess dynamic changes in ISF levels of Aβ, glucose, and lactate during hyperglycemia in Kir6.2+/+, APP/PS1 and Kir6.2-/-, APP/PS1 mice. (B) No difference in body weights or fasted blood glucose levels in APP/PS1 mice with or without Kir6.2-KATP channels prior to hyperglycemic clamp. (C) Kir6.2-/-, APP/PS1 mice are glucose intolerant. (D) While no difference in fasting insulin levels existed at baseline, hyperglycemia increased insulin levels in the Kir6.2+/+, APP/PS1 mice 2.1-fold while insulin levels did not change in Kir6.2-/-,APP/PS1 mice. Blood glucose levels at baseline and during the hyperglycemic clamp did not differ between groups. Hyperglycemia caused a 1.1-fold and 1.5-fold increase in blood glucose levels in Kir6.2+/+ or Kir6.2-/-, APP/PS1 mice, respectively. (E) Blood glucose levels were higher during the first 2 hours of the clamp in Kir6.2-/-, APP/PS1 mice compared to Kir6.2+/+, APP/PS1 mice. (R) There was a 3.2-fold decrease in the glucose infusion rate for the Kir6.2-/-, APP/PS1 mice compared to Kir6.2+/+, APP/PS1 mice due to an attenuated insulin response.

At baseline, there was no difference in body weight, blood glucose, or fasting plasma insulin levels in Kir6.2+/+, APP/PS1 and Kir6.2-/-, APP/PS1 mice (Figure 2B&D). However, the Kir6.2-/-, APP/PS1 mice are glucose intolerant compared to Kir6.2+/+, APP/PS1 mice (Figure 2C). The target blood glucose range for hyperglycemic clamps was 250-300mg/dL, resulting in a 100-150% increase, or doubling, of blood glucose levels relative to fasted baseline in Kir6.2+/+, APP/PS1 (p < 0.0001) and Kir6.2-/-, APP/PS1 mice (p < 0.0001; Figure 2D). In response to hyperglycemia, Kir6.2+/+, APP/PS1 mice also showed a 210% increase in plasma insulin compared to fasting levels (p = 0.002), while no change on plasma insulin levels were observed in Kir6.2-/-, APP/PS1 mice (p = 0.8976). These results confirm previous findings that glucose induced insulin secretion is impaired in Kir6.2-/- mice [31]. Reduced insulin secretion in Kir6.2-/-, APP/PS1 mice was also accompanied by higher blood glucose levels during the first 2 hours of the clamp (Figure 2E) and a 68% reduction in glucose infusion rates compared to Kir6.2+/+, APP/PS1 mice (p = 0.0003; Figure 2F). Regardless, blood glucose levels in Kir6.2+/+, APP/PS1 and Kir6.2-/-, APP/PS1 mice were comparable during the 4-hour hyperglycemic clamp despite the differential effect on plasma insulin levels and the insulin response.

When blood glucose doubled during the hyperglycemic clamp, hippocampal ISF glucose levels increased by 78±11% and 96±18% in Kir6.2+/+, APP/PS1 and Kir6.2-/-, APP/PS1 mice, respectively (Figure 3A). In Kir6.2+/+, APP/PS1 mice, increased ISF glucose levels were associated with a 19±6.5% increase in ISF Aβ (Figure 3B). Conversely, no hyperglycemia-dependent increase in ISF Aβ levels was observed in Kir6.2-/-, APP/PS1 mice (Figure 3B). ISF lactate is the metabolic end product of glycolysis and used as a marker of neuronal activity, where it covaries with increased EEG amplitude and ISF Aβ levels [9, 12, 14, 26, 39]. In Kir6.2+/+, APP/PS1 mice, ISF lactate increased by ∼22±8.4% during hyperglycemia, while no increase in ISF lactate was observed in Kir6.2-/-, APP/PS1 mice. Together, these data demonstrate that Kir6.2-KATP channels are necessary to couple changes in ISF glucose with ISF Aβ and ISF lactate.

**Figure 3.**
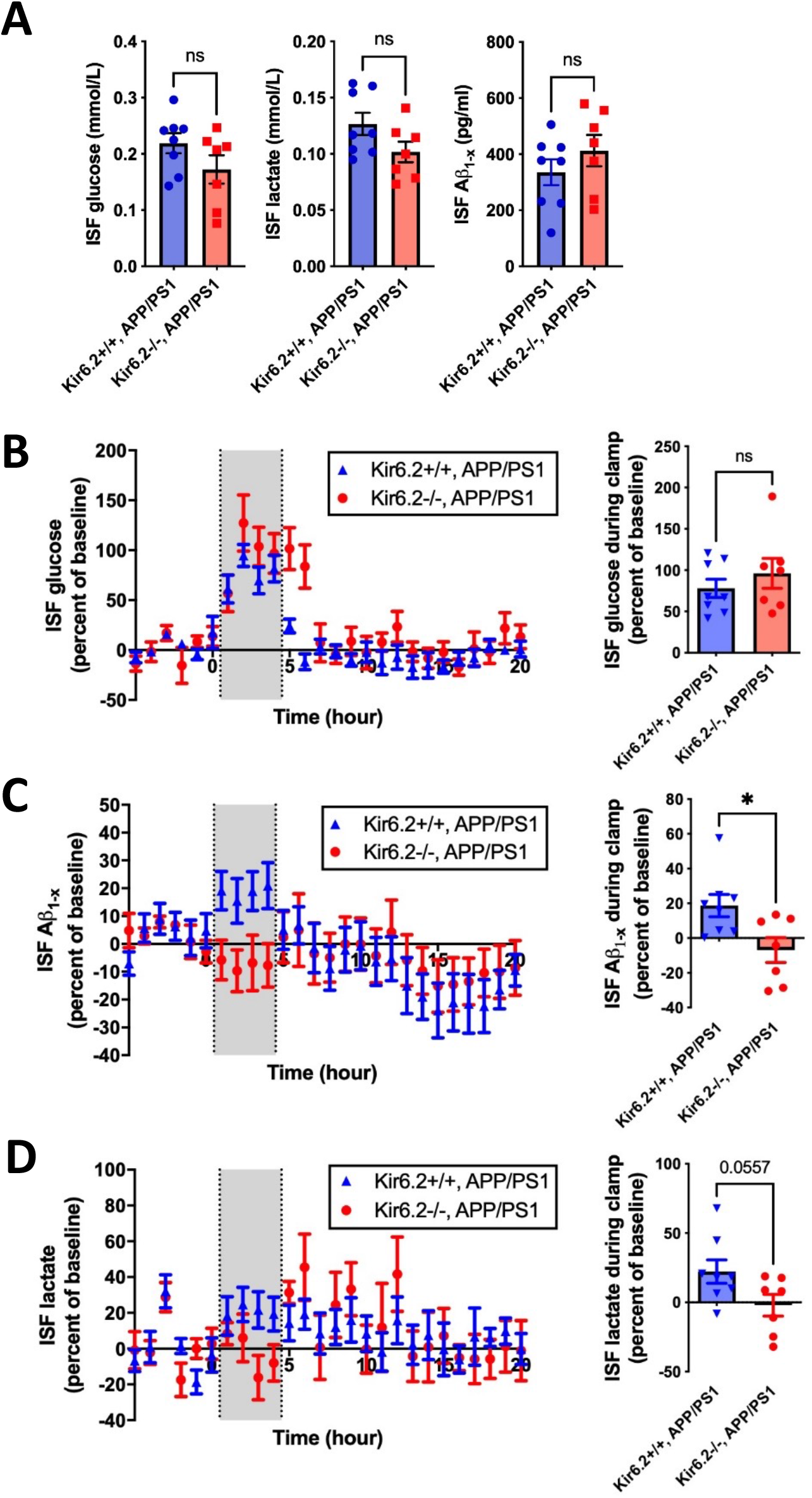
The effects of hyperglycemic clamps on ISF glucose, ISF Aβ, and ISF lactate in Kir6.2-/-, APP/PS1 and Kir6.2+/+, APP/PS1 mice. (A) Steady state levels of ISF glucose, lactate, and Aβ were similar between APP/PS1 mice with and without Kir6.2-KATP channels. (B) ISF glucose levels increased during the clamp to comparable levels in Kir6.2-/-, APP/PS1 and Kir6.2+/+, APP/PS1 mice (n=7-8 mice/group). (C) Although ISF glucose levels were comparable between groups, ISF Aβ increased by 19% in Kir6.2+/+, APP/PS1 mice, while no increase in ISF Aβ was observed in Kir6.2-/-, APP/PS1 mice. (D) Similarly, ISF lactate increased in Kir6.2+/+, APP/PS1 mice by 22% while no increased was observed in Kir6.2-/-, APP/PS1 mice.

Pearson’s correlations were performed on hippocampal ISF glucose, lactate, and Aβ levels during hyperglycemic clamp to further explore the effects of KATP channel deletion on these relationships. ISF Aβ levels showed strong, positive correlations (Figure 4A) with glucose (r = 0.5501, p = 0.0011) and lactate (r = 0.8079, p < 0.0001) in Kir6.2+/+, APP/PS1 mice. Similarly, ISF glucose and lactate also displayed a strong, positive correlation in Kir6.2+/+, APP/PS1 mice (r = 0.6875, p < 0.0001), demonstrating a positive relationship between energy demand, neuronal activity, and Aβ release. The relationships between ISF glucose, Aβ, and lactate were uncoupled Kir6.2-/-, APP/PS1 mice. The correlation between ISF glucose and ISF Aβ as well as ISF glucose and ISF lactate were lost in Kir6.2-/-, APP/PS1 mice (Figure 4B), while a correlation between ISF Aβ and lactate (r = 0.5268, p = 0.0048) persisted but only when ISF Aβ decreased (Figure 4B). This suggests that 1) KATP channels couple ISF glucose and ISF lactate and 2) ISF lactate may be necessary for increases in ISF Aβ. Regardless, Kir6.2-KATP channels are necessary for the hyperglycemia-dependent increases in hippocampal ISF Aβ and ISF lactate.

**Figure 4.**
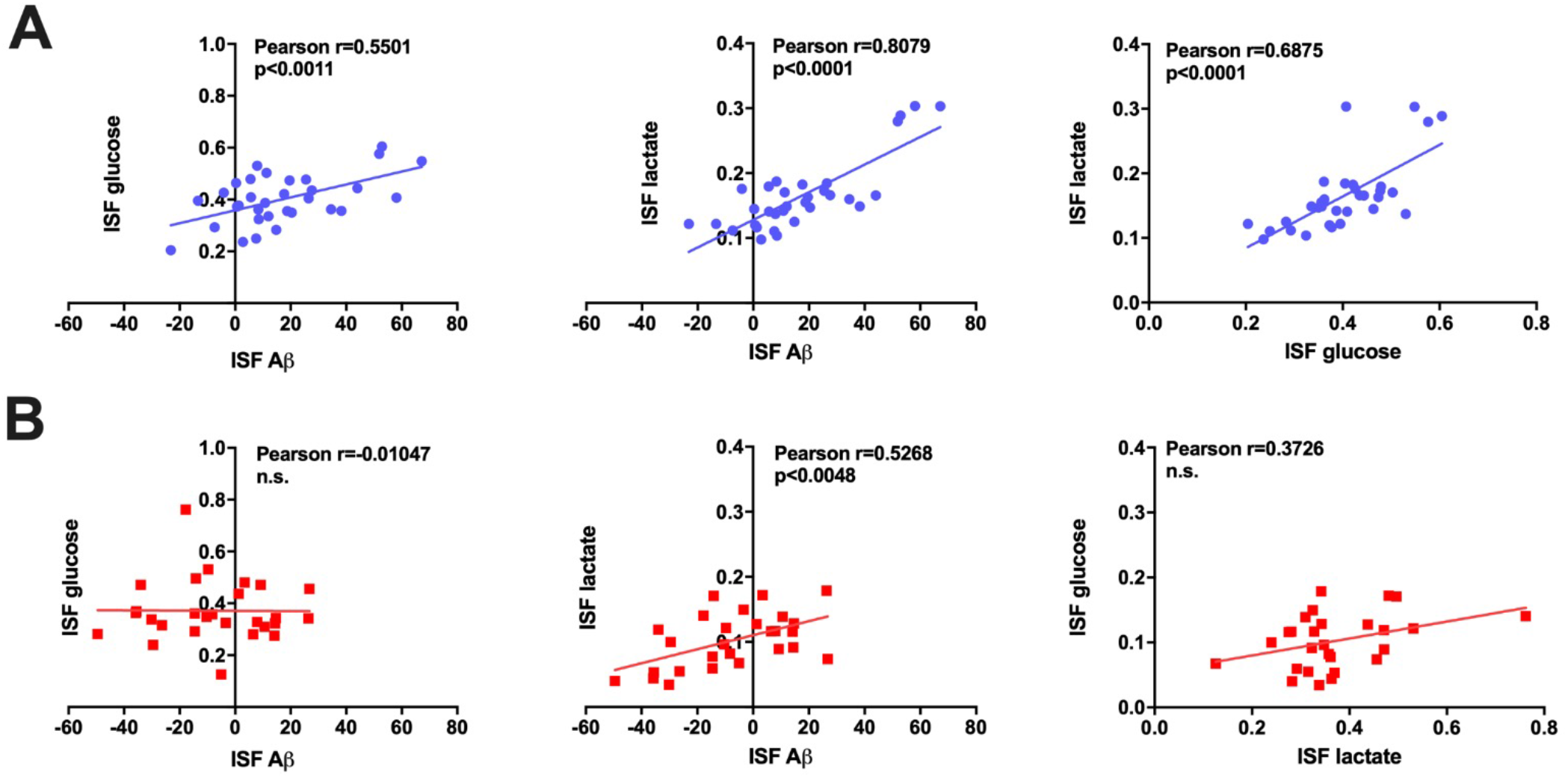
Correlations between ISF glucose, ISF Aβ, and ISF lactate in APP/PS1 mice with or without Kir6.2-K_ATP_ channels during hyperglycemic clamps. (A) In Kir6.2+/+, APP/PS1 mice (blue), ISF glucose, ISF Aβ, and ISF lactate display a positive correlation. (B) Conversely, in Kir6.2-/-, APP/PS1 mice (red), ISF glucose is no longer correlated with ISF Aβ or ISF lactate.

To further clarify the role of K_ATP_ channels in glucose-dependent increases in ISF Aβ, we explored whether inhibition of Kir6.1-K_ATP_ channels, those predominantly localized to the vasculature, also increased ISF Aβ levels. APP/PS1 mice deficient in Kir6.1 (Kir6.1-/-, APP/PS1) or Kir6.2 (Kir6.2-/-, APP/PS1) received a hippocampal infusion of KATP channel antagonist, glibenclamide, via reverse microdialysis. Interestingly, glibenclamide infusions raised ISF Aβ levels in both Kir6.2+/+, APP/PS1 and Kir6.1-/-, APP/PS1 mice by 29±9.1% and 24±6.7%, respectively, but not in APP/PS1, Kir6.2-/- mice (Figure 5A&B). This suggests that Kir6.2-KATP channels are responsible for the changes in ISF Aβ during hyperglycemia, not Kir6.1-KATP channels. Since Kir6.1-KATP channels are comprised of different subunits and localized to the vasculature, this suggests that neuronal KATP channel activity controls glucose-dependent increases in ISF Aβ.

**Figure 5.**
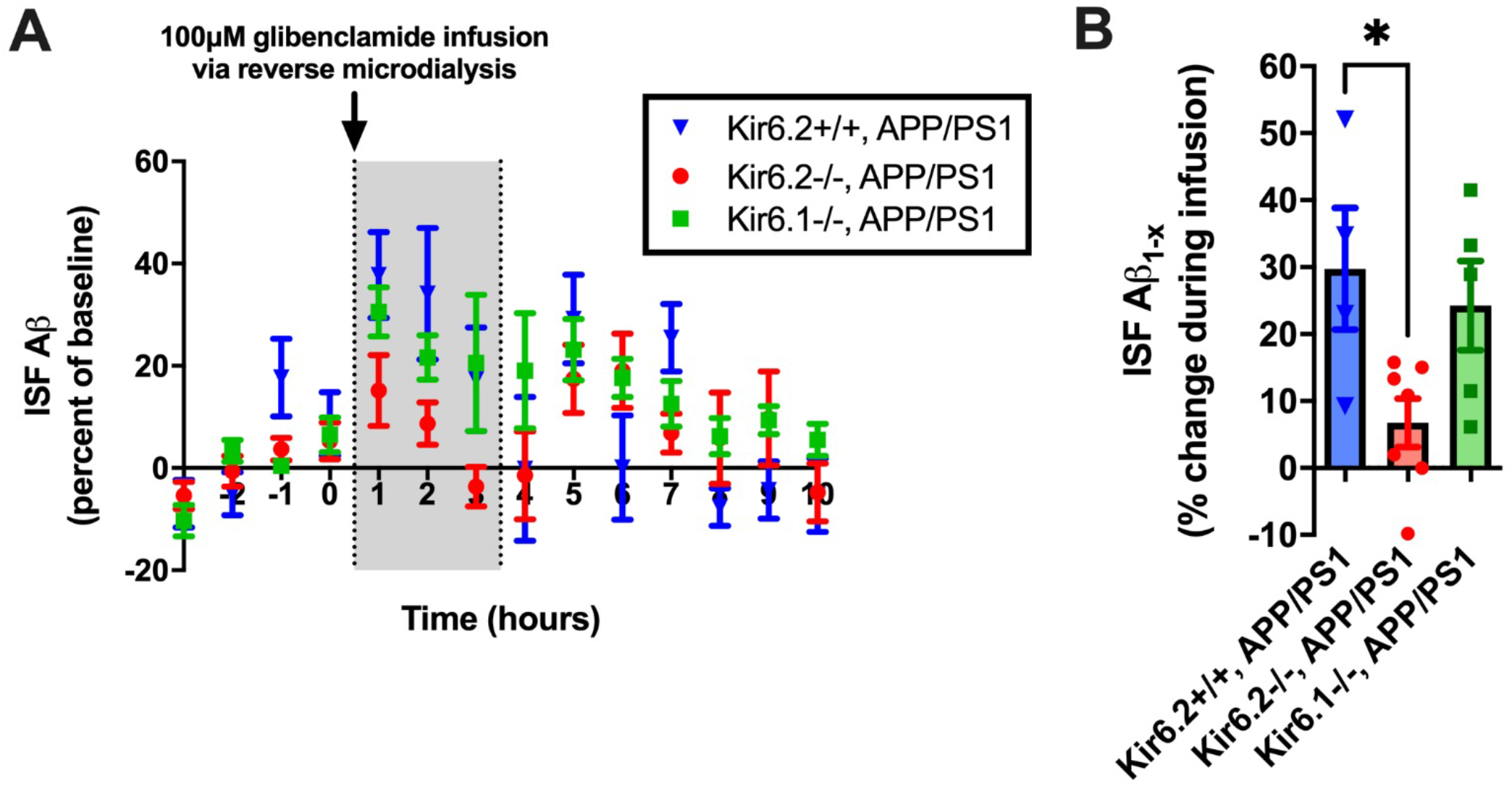
Glibenclamide infusion via reverse microdialysis modulates ISF Aβ levels in Kir6.2+/+, APP/PS1 and Kir6.1-/-, APP/PS1 mice but not Kir6.2-/-, APP/PS1 mice. (A) 100μm glibenclamide was infused directly into the hippocampus of Kir6.2+/+, APP/PS1 (blue triangles), Kir6.1-/-, APP/PS1 (red circles), or Kir6.2-/-, APP/PS1 (green squares) mice (n=4-7 mice/group). ISF Aβ levels were measured hourly during the 4-hr baseline, 3-hr infusion, and 6-hrs post-infusion. (B) During the glibenclamide infusion, ISF Aβ levels rose in Kir6.2+/+,APP/PS1 mice or Kir6.1-/-, APP/PS1 mice by 30% or 24%, respectively. Conversely, ISF Aβ levels rose by only 6% in Kir6.2-/-, APP/PS1 mice which was different than both other groups, suggesting Kir6.2-containing K_ATP_ channels are necessary for KATP channel dependent increases in ISF Aβ.

### Kir6.2-KATP channels couple chronic changes in peripheral glucose levels with Aβ pathology

Beginning at 3 months of age, Kir6.2+/+, APP/PS1 and Kir6.2-/-, APP/PS1 mice were exposed to high glucose, high fructose drinking water (e.g. sucrose water) or normal drinking water for 6 months to determine whether Kir6.2-KATP channels are necessary for glucose-dependent increases in amyloid plaque formation and AD-related pathology (n=8-11 mice/group; Figure 6A). Prior to sacrifice, body weight and blood glucose levels were collected 9 months of age. Terminal plasma insulin, glucose, and lactate measures were also taken to determine the effects of high sucrose consumption on peripheral metabolism to determine how it related to changes in on AD-related pathology within the brain.

**Figure 6.**
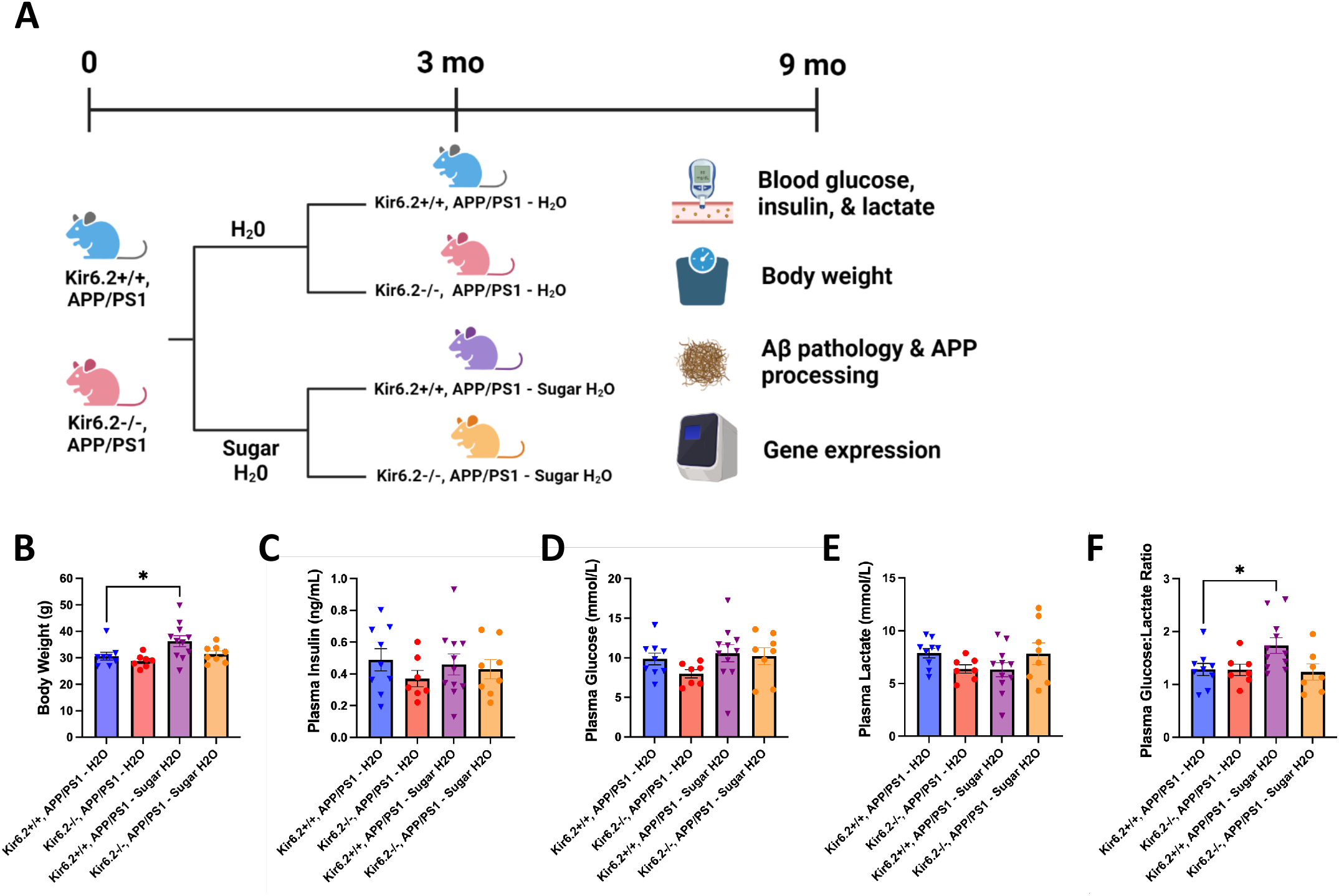
Chronic sugar exposure differentially effects peripheral metabolism in Kir6.2+/-, APP/PS1 and Kir6.2-/-, APP/PS1 mice. **(A)** Schematic of experimental approach where Kir6.2+/+, APP/PS1 and Kir6.2-/-, APP/PS1 mice were fed regular drinking water or high glucose, high fructose drinking water for 6 months. **(B)** Kir6.2+/+, APP/PS1 – Sugar H_2_O mice showed increased body weight compared to Kir6.2+/+, APP/PS1 – H_2_O mice (p = 0.0443). There were no changes in Kir6.2-/-, APP/PS1 mice fed regular drinking water or sugar drinking water. **(C)** Plasma insulin levels, **(D** plasma glucose levels, or **(E)** plasma lactate levels were comparable between all groups after 6 months on regular or sugar water. **(F)** Kir6.2+/+, APP/PS1 – Sugar H_2_O showed an increase the plasma glucose: lactate ratio compared to Kir6.2+/+, APP/PS1 – H_2_O mice (p = 0.0459).

At 9 months of age, Kir6.2+/+, APP/PS1 mice who consumed sucrose H_2_O for 6 months weighed ∼20% more than Kir6.2+/+, APP/PS1 fed normal tap water (p = 0.0443; Figure 6B). Interestingly, neither high sucrose consumption nor genotype had any effect on plasma glucose, plasma insulin, or plasma lactate levels in any group assayed at a terminal time point (Figure 6C-E). Interestingly, Kir6.2+/+, APP/PS1 – sucrose H_2_O mice had a 35% increase in the ratio of plasma glucose:lactate compared to Kir6.2+/+, APP/PS1 – H_2_O mice (p = 0.0459), suggesting high levels of glucose were present in the blood.

To evaluate how high sucrose diet affected Aβ-related pathology in Kir6.2+/+, APP/PS1 and Kir6.2-/-, APP/PS1 mice, we quantified Aβ deposition via immunohistochemistry and amyloid precursor protein (APP) processing via western blot analysis (Figure 7). First, 50 μm sections were immunostained with the monoclonal Aβ antibody, HJ3.4B, to identify Aβ plaques within the cortex and hippocampus. Percent area coverage in both of these regions was quantified for the following 4 groups: 1) Kir6.2+/+, APP/PS1 - H_2_O mice, 2) Kir6.2-/-, APP/PS1 mice - H_2_O, 3) Kir6.2+/+, APP/PS1 – sucrose H_2_O mice, and 4) Kir6.2-/-, APP/PS1 – sucrose H_2_O mice (Figure 7A). Genotype alone did not impact Aβ deposition in the APP/PS1 mice. Kir6.2+/+, APP/PS1 mice and Kir6.2-/-, APP/PS1 mice fed normal drinking had comparable levels of Aβ deposition in both cortex and hippocampus. Conversely, Kir6.2+/+, APP/PS1 mice fed a high sucrose diet had 2-3 times more Aβ plaques in the cortex (p = 0.0262) and hippocampus (p = 0.0202) compared to Kir6.2+/+, APP/PS1 mice fed a normal diet (Figure 7B). This suggests that chronic overconsumption of sucrose is sufficient to increase Aβ deposition in mice. Interestingly, mice lacking Kir6-2-KATP channels fed a high sucrose diet did not display the same increase in Aβ deposition as Kir6.2+/+, APP/PS mice (Figure 7A). This suggest that Kir6.2-KATP channels are necessary for sugar-dependent increases in Aβ pathology in mice.

**Figure 7.**
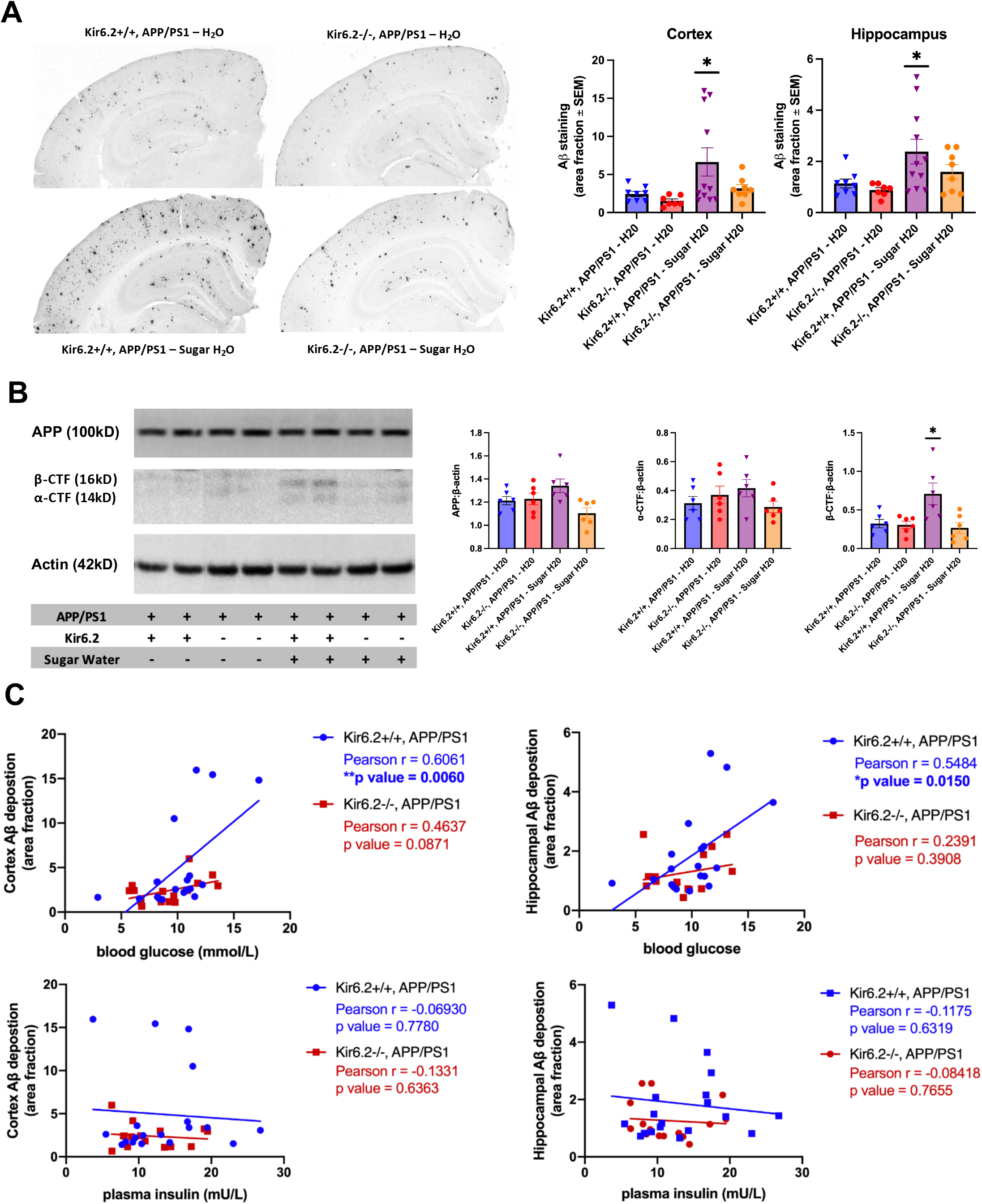
APP/PS1 mice fed a high sugar diet had increased Aβ deposition and amyloidogenic process of amyloid precursor protein (APP). **(A)** Representative images of Aβ deposition in Kir6.2+/+, APP/PS1 and Kir6.2-/-, APP/PS1 mice on fed a normal or high sugar diet. Aβ deposition was increased in both the cortex (p = 0.0262) and the hippocampus (p = 0.0202) of the Kir6.2+/+, APP/PS1 mice fed a high sugar diet but not in Kir6.2-/-, APP/PS1 mice fed a high sugar diet. **(B)** Western blot analysis for APP and the C-terminal fragments (CTF) showed no difference in the expression of full-length APP or α-CTF between any groups, but there was an increase in β-CTF in Kir6.2+/+, APP/PS1 mice fed a high sugar diet. **(C)** In kir6.2+/+, APP/PS1 mice, there is a significant, positive correlation between blood glucose levels and Aβ deposition in both the cortex (Pearson r=0.6061, p=0.006) and hippocampus when mice were on a regular or high sugar diet (Pearson r =0.5485, p=0.0150). Conversely, in Kir6.2-/-, APP/PS1 mice, no correlation exists between blood glucose levels and Aβ deposition in either the cortex or hippocampus. Plasma insulin levels were not correlated with Aβ deposition in any condition.

To determine whether chronic sucrose consumption altered APP metabolism, APP processing was analyzed through measurement of total APP as a well as the protein levels of two key APP metabolites, CTF-α and CTF-β (Figure 7B). There were no changes in the levels of APP or CTF-α due to genotype or sucrose H_2_O (Figure 7A). Importantly, there was increased CTF-β in Kir6.2+/+, APP/PS1 – sucrose H_2_O compared to all other groups (p = 0.0052), suggesting that chronic sucrose consumption increases the amyloidogenic processing of APP to generate the cleavage productions, CTF-β and Aβ. Together, these data suggest that chronic exposure to sucrose H_2_O, Kir6.2 containing K_ATP_ channels increased both Aβ deposition (Figure 7A) and amyloidogenic processing of APP production (Figure 7B) in the brains of Kir6.2+/+, APP/PS1 mice; an effect that was lost with Kir6.2-KATP channel deletion.

Next, we explored how changes in peripheral metabolism correlated with changes in Aβ-related pathology in Kir6.2+/+, APP/PS1 and Kir6.2-/-, APP/PS1 mice (Figure 7C). In Kir6.2+/+, APP/PS1 mice, peripheral blood glucose levels correlated with Aβ deposition in both the cortex (Pearson r=0.6061, p value =0.0060) and hippocampus (Pearson r=0.5484, p value=0.0150). In APP/PS1 mice lacking Kir6.2, no significant correlations persisted between blood glucose level and Aβ pathology. We also explored whether plasma insulin, another metabolite implicated in AD pathogenesis, correlated with Aβ pathology; however, no correlation between insulin and Aβ existed in either genotype or diet. This suggests that increased blood glucose levels are sufficient to drive Aβ pathology and Kir6.2-KATP channels are necessary to do so.

### KATP channel activity is necessary for hyperglycemia-dependent release of astrocytic lactate

Our data suggest that KATP channels are necessary for glucose-dependent increases in lactate and Aβ. Therefore, we explored how deletion of KATP channels impacted the metabolism and transport of glucose and lactate. According to the astrocyte neuron lactate shuttle (ANLS), glucose uptake, lactate production, and lactate transport are compartmentalized in astrocytes and neurons [42]. Astrocytes take up glucose from the blood stream via GLUT1 and process it glycolytically via HK1 and GAPDH. Pyruvate is then converted to lactate via lactate dehydrogenase A (LDHA) and released extracellularly via monocarboxylate transporter 4 (MCT4). Lactate uptake into neurons is mediated by monocarboxylate transporter 2 (MCT2), where it is converted back to pyruvate via lactate dehydrogenase B (LDHB) for use in mitochondrial respiration and oxidative phosphorylation via pyruvate dehydrogenase (PDH; Figure 8A). Therefore, we explored whether Aβ pathology, KATP channel activity, and high sucrose diet affected the expression of *Glut1, Glut3, Ldha, Ldhb, Mct2, Mct4, Hk1, Gaphd, and Pdha1* in APP/PS1 mice, +/- Kir6.2-KATP channel expression and +/- high sucrose diet. Comparing genotype alone, Kir6.2+/+, APP/PS1 mice displayed decreased *Pdha1, Ldha*, and *Ldhb* expression compared to WT mice; an effect not observed in Kir6.2-/-, APP/PS1 mice (Figure 8B). In mice fed a high sugar diet, Kir6.2+/+, APP/PS1 mice had decreased *Glut1* and *Mct2* expression compared to WT, suggesting decrease glucose uptake in astrocytes and decrease lactate uptake in neurons, while displaying increased *Hk1* expression, suggesting increased glycolysis. On the other hand, Kir6.2-/-, APP/PS1 on a high sugar diet seemed to have an opposing phenotype, where *Mct2* expression increased, *Pdha1* increased, and *Mct4* decreased, suggesting less astrocytic lactate release, conserved neuronal lactate uptake, and preserved mitochondrial respiration. Together, this suggests that astrocytic release of lactate is modulated by Kir6.2-KATP channels and is important for Aβ release and aggregation.

**Figure 8.**
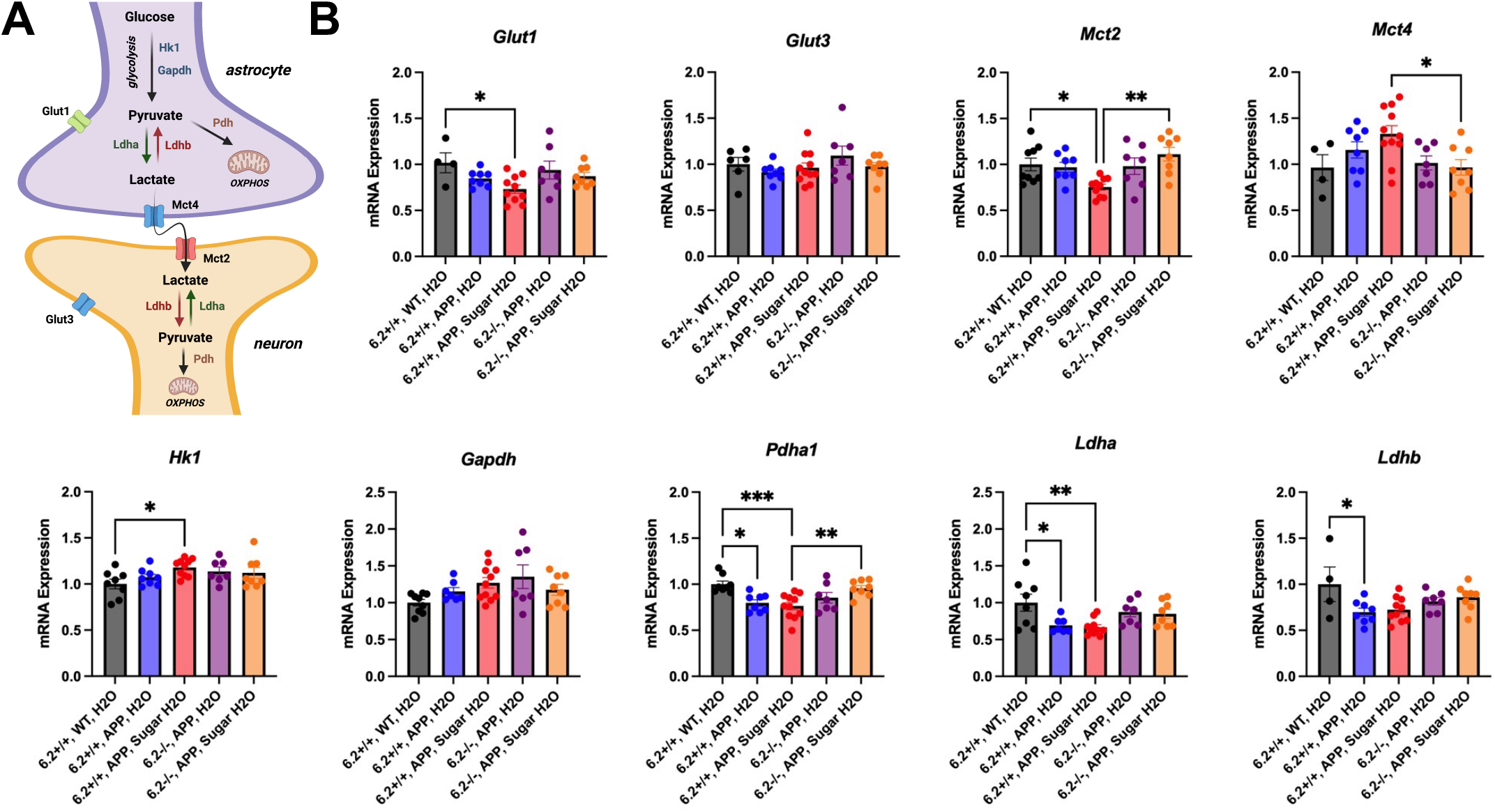
Alterations in lactate metabolism and transport due to the interaction between Aβ, K_ATP_ channels, and a high sucrose diet. **(A)** Simplified schematic of glucose uptake, lactate production, and lactate transport is compartmentalized in astrocytes and neurons. Astrocytes take up glucose via Glut1 and process it glycolytically (e.g. Hk1 and Gapdh). Pyruvate is converted to lactate via Ldha and released extracellularly via Mct4. Lactate uptake into neurons is mediated by Mct2, where it is converted back to pyruvate via Ldhb for use in mitochondrial respiration and oxidative phosphorylation. **(B)** Kir6.2+/+, APP/PS1 mice on a high sucrose diet had decreased *Glut1* and *Mct2* expression compared to WT, suggesting decrease glucose uptake in astrocytes and decrease lactate uptake in neurons, while displaying increased *Hk1* expression, suggesting increased glycolysis. Kir6.2-/-, APP/PS1 on a high sugar diet seem to have an opposing phenotype, where *Mct2* expression increased, *Pdha1* increased, and *Mct4* decreased, suggesting less astrocytic lactate release, conserved neuronal lactate uptake, and preserved mitochondrial respiration. Kir6.2+/+, APP/PS1 mice alone displayed decreased *Pdha1, Ldha*, and *Ldhb* expression, which was not mirrored in Kir6.2-/-, APP/PS1 mice.

## DISCUSSION

K_ATP_ channels modulate a wide array of physiological functions and their function has been meticulously detailed in pancreatic insulin release [43], cerebral metabolism [44-46], cardiovascular regulation [47-49], and neuronal excitability and Aβ release [50]. This study builds upon our previous hypothesis that hyperglycemia directly influences neuronal Aβ release and offers a mechanism through which hyperglycemia results in increased Aβ production and deposition. To begin, we confirm previous findings that CNS hyperglycemia results in Aβ release and assert that these effects are mediated specifically through Kir6.2 containing K_ATP_ channels.

The goal of these experiments was to expand upon our previous data that showed acute hyperglycemia is sufficient to cause increased Aβ release [50] and to elucidate the role of Kir6.2 containing K_ATP_ channels in this process. Under acute hyperglycemia, Kir6.2+/+, APP/PS1 mice released more Aβ into the hippocampal ISF compared to Kir6.2-/-, APP/PS1 demonstrating a relationship between Kir6.2, ISF glucose, and Aβ release. Regression analysis of ISF Aβ levels during hyperglycemia showed strong positive correlations between ISF Aβ, glucose, and lactate levels, affirming a metabolic influence on Aβ release [50-54]. Deletion of neuronal KATP channel activity abolished these relationships demonstrating KATP channels link changes in cerebral metabolism with Aβ production, release, and aggregation.

To further analyze this relationship, we utilized standard APP/PS1 mice with intact K_ATP_ channels, as well APP/PS1 mice lacking either Kir6.2 (primarily expressed on neurons) or Kir6.1 (primarily expressed on pericytes and endothelial cells) treated with a nonspecific K_ATP_ channel antagonist, glibenclamide. Glibenclamide works to reduce K_ATP_ channel dependent K^+^ efflux, resulting in a higher resting membrane potential, making affected cells more excitable [43, 45, 49, 55]. Since Aβ is released from neurons in an activity-dependent manner, cells that are more readily excited will release more Aβ [50, 56-58]. As such, both Kir6.2+/+, APP/PS1 and Kir6.1-/-, APP/PS1 mice treated with glibenclamide released significantly more Aβ than Kir6.2-/-, APP/PS1 mice. These data demonstrate, specifically, that Kir6.2 containing K_ATP_ channels facilitate Aβ release from neurons, not Kir6.1 containing K_ATP_ channels. These data, taken together with the hyperglycemic clamp data, clearly demonstrate that **1)** Kir6.2 containing K_ATP_ channels are involved in activity dependent Aβ release and **2)** hyperglycemia increases ISF Aβ release through Kir6.2-K_ATP_ channel modulation.

Interestingly, Kir6.2-/-, APP/PS1 showed no metabolic abnormalities or alterations in AD-related pathology at baseline, suggesting that dietary intervention is necessary to detect deficiencies in Kir6.2-/- mice. These data coincide with previous reports on the Kir6.2-/- phenotype, where Kir6.2-/- alone is not sufficient to cause overt changes in fasting blood glucose or insulin secretion. In fact, a glucose challenge was necessary to observe an impairment in glucose sensing and subsequent buffering via insulin secretion [55]. However, Kir6.2-/-, APP/PS1 mice showed an impaired ability to release insulin in response to glucose incursion, which in turn, caused a longer time to buffer blood glucose levels during acute hyperglycemia (Figure 2). It is also interesting that Aβ deposition in Kir6.2+/+, APP/PS1 and Kir6.2-/-, APP/PS1 mice did not differ at baseline, only in response to a chronic sucrose dietary challenge. First, it could be argued that neuronal KATP channel deletion should increase the resting membrane potential leading to chronic hyperexcitability. This, in turn, would increase Aβ production and amyloid plaque formation. However, we did not observe any differences in Aβ deposition in Kir6.2+/+, APP/PS1 and Kir6.2-/-, APP/PS1 mice consuming regular drinking water, only in response to a hyperglycemic challenge. This suggests there is still more to learn about the role of neuronal KATP channel activity in AD.

Furthermore, we found no evidence of impaired glucose transport into the brain between Kir6.2+/+, APP/PS1 and Kir6.2-/-, APP/PS1 mice (Figure 2A). This is important for two reasons. First, it demonstrates that Kir6.2-KATP channels are not necessary glucose transport into the brain. These data coincide with transcriptomic data that show Kir6.2 (KCNJ11/*Kcnj11)* is predominantly expression on neurons, and only minimally expressed on astrocytes, pericytes, and endothelial cells [59]. Since endothelial cells, pericytes, and astrocytes are responsible for the initial uptake of glucose from the blood to brain [60], it is unlikely that the absence of Kir6.2- K_ATP_ channels would affect energy availability in the brain. Similarly, these studies reinforce the idea that glucose uptake into the brain is largely insulin independent. Since comparable levels of ISF glucose were attained in both the Kir6.2+/+, APP/PS1 and Kir6.2-/-, APP/PS1 brains with a hyperglycemia clamp, but the Kir6.2-/-, APP/PS1 mice did not have a compensatory increase in blood insulin levels, this demonstrates that glucose transport from the blood to brain is not dependent on nor altered by plasma insulin levels.

These experiments are instrumental in demonstrating a mechanistic relationship between Kir6.2-K_ATP_ channels, glucose metabolism, and Aβ release. Interestingly, our studies also demonstrate a unique relationship between KATP channel activity and lactate production, which may be a necessary step for extracellular Aβ release and subsequent aggregation. It is well documented that ISF and CSF lactate levels display a strong correlation with ISF and CSF levels of Aβ and tau, the pathological hallmarks of AD [12, 14, 26, 39, 61-63]. ISF lactate levels covary with ISF Aβ and ISF tau across the circadian day and sleep/wake cycles, where ISF Aβ, tau, and lactate increase during periods of sleep deprivation [14, 62]. During increased periods of neuronal activity, ISF lactate rises and is strongly correlated with increased ISF Aβ [39]. In the human brain, aerobic glycolysis, a process where excess glucose is used for lactate production not oxidative phosphorylation, is a biomarker of brain regions vulnerable to amyloid and tau deposition [64-66]. Thus, extracellular lactate levels, at minimum, covary with Aβ and tau, or more likely, plays a role in Aβ and tau release. Our data suggest that KATP channels couple glucose and lactate, representing an opportunity to intervene therapeutically with the potential of modulating Aβ and tau levels. Our studies also suggest that targeting lactate might be important for reducing Aβ, and perhaps tau, release and aggregation.

We also aimed to discern how a mild metabolic insult via chronic exposure to sucrose H_2_O affects AD-related pathology in APP/PS1 mice +/- Kir6.2-K_ATP_ channels. It is well established that T2D and metabolic syndrome (MetS) are strong risk factors for the development of AD [67-69], but this study reinforces the idea that a subtle change in glucose homeostasis, in the absence of T2D, MetS, or hyperinsulinemia, is sufficient to alter APP processing, Aβ production and AD-related pathology [50, 54, 70-73]. It is important to note that this paradigm was not intended to bring about an over diabetic or obese phenotype, but rather to analyze how a specific change in blood glucose levels over time might alter the delicate balance of cerebral metabolism and AD-related pathology.

In the present study, we found that a mild sucrose H_2_O insult in animals with intact Kir6.2+/+ K_ATP_ channels had increased Aβ plaque deposition, ISF Aβ levels, and CTF-β expression. These data offer a stepwise mechanism to demonstrate how Kir6.2-K_ATP_ channels facilitate increases in Aβ plaque deposition: 1) Hyperglycemia causes aberrant metabolism that is mediated through the presence of Kir6.2 K_ATP_ channels. 2) This change in neuronal metabolism causes the amyloidogenic processing of APP, resulting in higher CTF-β and Aβ generation [74]. 3) Increased stimulation of hippocampal neurons via hyperglycemia and Kir6.2 K_ATP_ channels results in increased Aβ release into ISF. 4) Over time, increased concentration of extracellular Aβ aggregates into extracellular amyloid plaques in a concentration dependent manner.

Lastly, these studies demonstrate a novel mechanism by which individuals with prediabetes or type-2-diabetes may be at an increased risk to develop AD. It demonstrates that increased sucrose intake or elevations in blood sucrose are sufficient to drive Aβ pathology, independent of T2D, obesity, and MetS. Importantly, it also suggests that pharmacological antagonism of Kir6.2-K_ATP_ channels holds therapeutic promise for diabetic and prediabetic patients to help reduce risk of developing AD.

## CONCLUSIONS

Though many human studies have linked obesity, metabolic syndrome, and diabetes to an increased likelihood of developing AD [67, 75-80] and numerous rodent studies have shown a link between diabetes and AD [53, 54, 71-73, 81-83], few studies have utilized a moderate metabolic challenge to isolate the effects of hyperglycemia and glucose dyshomeostasis on the development of AD. In acute and chronic paradigms, exposure to hyperglycemia or sucrose H_2_O, respectively, caused an increase in ISF Aβ levels, amyloidogenic processing of APP, and Aβ pathology, which was mediated by Kir6.2 containing K_ATP_ channel activity.

Importantly, this study shows that Kir6.2+/+ containing K_ATP_ channels couple changes in glucose metabolism with neuronal excitability and Aβ metabolism.

## AUTHOR CONTRIBUTIONS

SLM, DMH, and MP conceived of the study. SLM, DMH, MSR, CGN, JG, and MP contributed to study design. SLM, JG, MSS, EEC, WRM, CMC, JAS, SK, LD, NN, and DK performed experiments, data analysis, and data interpretation. JG and SLM wrote the manuscript. All authors discussed the results and commented on the manuscript.

## ACKNOWLEDGEMENTS

We would like to acknowledge the following grants: 1K01AG050719 (SLM), R01AG068330 (SLM), BrightFocus Foundation (A20201775S; SLM), Charleston Conference on Alzheimer’s disease New Vision Award (SLM), the McDonnell Center for Systems Neuroscience (SLM and AQB), P01NS080675 (DH, SLM), R01 HL140024 (CGN), F31AG066302 (CC), T32AG033534 (JG), F31AG071119 (MP) and T32AA007565 (SD).

